# Direct Prediction of the Complex-Valued Analytic Signal of EEG from Raw Multichannel Data

**DOI:** 10.1101/2024.05.13.593119

**Authors:** Takayuki Onojima, Takashi Imai

## Abstract

Accurate estimation of instantaneous neural dynamics is essential for electroencephalography (EEG)-based brain–state analysis and future closed-loop applications. Conventional phase estimation methods that rely on bandpass filtering and autoregressive prediction are limited by reduced accuracy near the current time point, poor future prediction capability, and susceptibility to inconsistencies in the contributing electrodes. This study proposes a deep neural network (DNN) framework that directly predicts the complex-valued analytic signal of EEG—representing instantaneous phase and amplitude—from raw multichannel EEG data. The model employs a temporal convolutional encoder and a probabilistic output head that jointly estimate the mean and variance of analytic signals at multiple past and future time points. To enhance robustness, a missing-signal imitation (MSI) mechanism is applied during training, together with electrode-mixing strategies incorporated into the model. Using resting-state EEG from 25 participants, we show that the proposed model consistently outperforms an autoregressive baseline in both subject-wise and cross-subject evaluations. MSI further improves stability when critical electrodes are removed and yields better calibration of predictive uncertainty. These findings demonstrate that analytic EEG signals can be estimated directly from raw data without manual preprocessing or subject-specific calibration. The proposed framework provides a scalable basis for automated EEG analysis and offers strong potential for future real-time and closed-loop neural applications.

## 1. Introduction

Estimating the instantaneous phase and amplitude of oscillatory components in electroencephalography (EEG) is essential for time–frequency analysis, artifact-free feature extraction, and real-time state tracking in signal processing applications. The analytic signal—obtained via the Hilbert transform—provides a principled representation of instantaneous oscillatory dynamics and is widely used in EEG analysis [1–3]. Conventional approaches typically combine bandpass filtering with autoregressive (AR) forward prediction to estimate instantaneous phase from ongoing EEG [4–6]. However, these methods suffer from reduced accuracy around the current time point due to edge effects in zero-phase filtering, limited capability for future prediction, and sensitivity to instability in the electrode signals. In addition, they rely on manually selected electrodes and frequency bands and operate primarily on single-channel signals or fixed spatial filters, thereby failing to exploit the spatial redundancy inherent in multichannel EEG recordings.

Recent advances in deep learning have motivated the development of end-to-end frameworks that learn neural representations directly from raw EEG [7–10]. Although these models have demonstrated improved within-subject performance, most remain sensitive to inter-individual variability and exhibit substantial degradation when applied to unseen subjects [7,8]. This limitation poses a significant challenge for EEG applications where participant-specific calibration is impractical or undesirable [7]. Addressing cross-subject generalization is therefore crucial for the broader adoption of predictive EEG models in large-scale experiments, passive monitoring systems, or plug-and-play neural decoding applications.

In this study, we introduce a deep neural network (DNN) architecture designed to directly predict analytic EEG signals—including both amplitude and phase—from raw multichannel EEG. The proposed model integrates a temporal convolutional encoder based on dilated causal convolutions [11], together with a probabilistic output head that estimates the mean and variance of the complex-valued analytic signal using a negative log-likelihood formulation [12,13]. To enhance robustness and reduce over-reliance on individual electrodes, we incorporate a missing-signal imitation (MSI) module that simulates channel dropout during training, as well as electrode-mixing strategies that include both learned cross-electrode integration and adjacency-guided graph-based aggregation [14]. These components mitigate the impact of channel variability and help stabilize prediction under realistic conditions where sensor dropout and spatial inconsistencies frequently arise.

Using resting-state EEG data from 25 participants, we demonstrate that the proposed method achieves high prediction accuracy not only within subjects but also in cross-subject evaluations, in which the model is tested on individuals not included during training. This ability to generalize across participants while preserving prediction fidelity represents an important step toward the development of scalable EEG prediction systems and lays the groundwork for future real-time and closed-loop applications.

In summary, the main contributions of this study are as follows: (1) we introduce a deep neural network that directly predicts complex-valued analytic EEG signals from raw multichannel input; (2) we incorporate a missing-signal imitation (MSI) mechanism and electrode-mixing strategies to enhance robustness against channel dropout; (3) we demonstrate improved prediction accuracy over an autoregressive baseline in both subject-wise and cross-subject evaluations; and (4) we show that MSI improves calibration of predictive uncertainty, enabling more reliable probabilistic predictions for future real-time applications.

## 2. Methods

### 2.1 Conventional methods

The conventional state-informed stimulation method estimates the instantaneous EEG phase by combining a zero-phase bandpass filter with an autoregressive (AR) forward prediction model [4–6]. This approach enables feedforward prediction, in which the current and future EEG phases are estimated solely from past EEG samples within a short time window. As illustrated in Figures 1a–d, the method requires manual selection of both the signal of interest and the frequency band before prediction can be performed. The prediction procedure consists of the following four steps: (1) selecting the target signal, (2) selecting the frequency band, (3) AR-based forward prediction, and (4) analytic signal and phase estimation.

**Figure 1:**
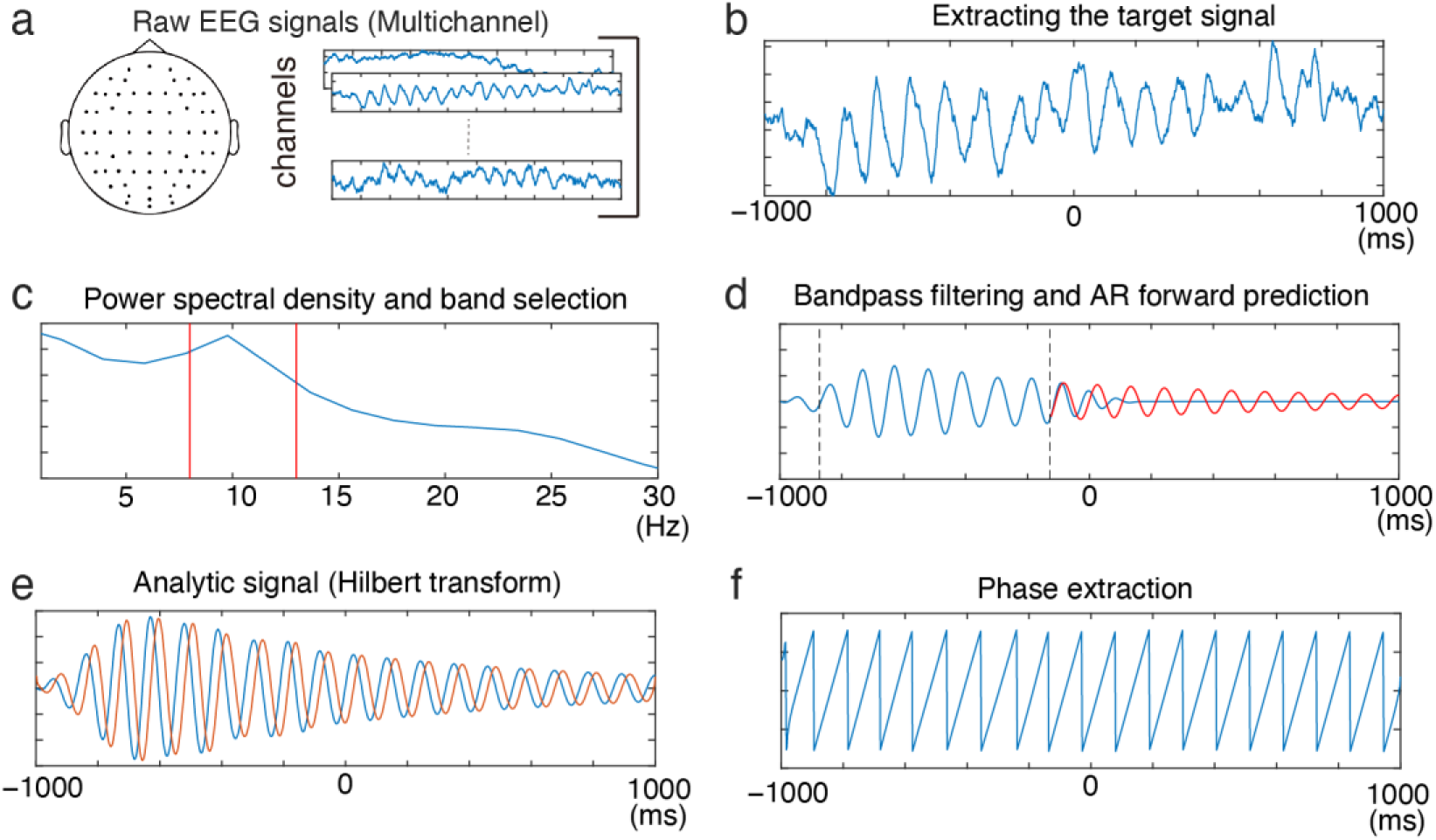
Conventional signal-processing pipeline for estimating instantaneous EEG phase. (a) Raw multichannel EEG signals. (b) Extracting the target signal, such as selecting an electrode or applying a spatial filter. (c) Power spectral density of the extracted signal and selection of the frequency band of interest. (d) Bandpass-filtered signal and its autoregressive (AR) forward prediction, where 0 ms denotes the current time in a real-time processing scenario. (e) Analytic signal obtained via the Hilbert transform applied to the predicted band-limited signal. (f) Instantaneous phase extracted from the analytic signal. In contrast to this multi-stage conventional pipeline, the proposed method directly predicts the analytic signal in (e) from the raw multichannel EEG in (a).

In the first step, the target signal is selected. EEG is recorded from multiple scalp electrodes (Figures 1a and 1b), and the electrode that best captures the oscillatory component of interest must be chosen manually, or alternatively, a spatial filter such as the Hjorth derivation may be applied to extract the signal of interest [15]. Because this approach processes each channel independently, it cannot exploit the spatial redundancy available in multichannel EEG recordings and is therefore sensitive to noise and channel variability.

In the second step, a power spectral density analysis is used to determine the dominant oscillatory component (Figure 1c). A zero-phase bandpass filter is then applied to isolate the selected frequency band. Since zero-phase filtering is inherently non-causal and relies on future samples, the edge portions of the filtered signal must be discarded to avoid artifacts introduced by zero padding, which degrades prediction accuracy near the current time point.

In the third step, the band-limited signal is modeled using an AR process, and future samples are predicted from past filtered values (Figure 1d). However, AR models assume a linear and stationary generative process, whereas EEG dynamics are highly nonlinear and nonstationary, making accurate future prediction fundamentally difficult.

In the final step, the instantaneous phase is obtained by applying the Hilbert transform [1–3] to the filtered or predicted signal (Figures 1e and f). For a band-limited signal 𝑠(𝑡) and its Hilbert transform 𝑠_𝐻_(𝑡), the analytic signal is defined as

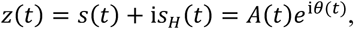

where 𝜃(𝑡) and 𝐴(𝑡) denote the instantaneous phase and amplitude, respectively.

In our experiments, this conventional pipeline also served as the baseline for comparison with the proposed DNN models. To ensure reproducibility, we briefly summarize the specific settings used for the AR-based method. For fair comparison, the same 8–13 Hz frequency band as the analytic-signal ground truth (Section 2.4) was used. A zero-phase FIR bandpass filter was applied via forward–reverse filtering (filtfilt in MATLAB), designed with the designfilt function (bandpassfir) at a sampling rate of 500 Hz and with a filter order of 128 taps (corresponding to a trimming window of 64 samples). The autoregressive model was estimated using the Yule–Walker method via the Levinson–Durbin recursion, with the model order set to 30, a commonly used configuration in real-time EEG phase estimation studies [4–6].

Although this conventional method enables real-time estimation of EEG phase, it has several limitations. Predicting future EEG samples is inherently difficult, and the use of a bandpass filter and AR model often results in suboptimal prediction accuracy. Furthermore, the need to manually select electrodes and frequency bands increases the complexity of the experimental setup and may introduce experimenter-dependent variability. These limitations motivate the development of an end-to-end approach that eliminates hand-crafted processing steps and directly predicts the analytic signal from raw EEG.

### 2.2 Deep neural network approach

To overcome the drawbacks of the conventional method, we employ a deep neural network (DNN) that directly predicts the analytic signal—comprising instantaneous phase and amplitude—from raw multichannel EEG, thereby bypassing the intermediate steps required in the conventional pipeline (i.e., the manual signal selection, bandpass filtering, and AR-based forward prediction in Figures 1b–d). In other words, the proposed model learns an end-to-end mapping from the raw EEG in Figure 1a to the analytic signal in Figure 1e.

#### 2.2.1 Input Representation

The network processes the most recent 1 s of EEG recorded from 63 electrodes. Let

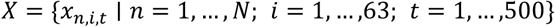

denote the input dataset, where 𝑁 is the number of samples, 63 is the number of electrodes, and 500 is the number of time points within each 1 s window. The model is trained to predict the current and future analytic signals for a selected frequency band.

#### 2.2.2 Temporal Encoder

The temporal encoder (Figure 2a) processes each input sample through three sequential stages: input standardization, a causal prefilter, and a dilated temporal convolutional network (TCN) stack [11], which generates temporal feature representations for each electrode. Standardization is applied per electrode along the time axis (zero mean, unit variance over time). The Prefilter is a causal one-dimensional convolution applied independently to each electrode, using a kernel length of 31 in our experiments and producing 4 output channels. The outputs are then normalized by layer normalization (applied over time and feature axes) and passed through a ReLU activation. Subsequently, a dilated TCN stack refines these representations via a sequence of causal temporal convolutions whose dilation factors grow exponentially, 𝑑 ∈ {1,2,4,8, … } , using the first 𝑛_blocks_ values. Each TCN block consists of a causal one-dimensional convolution with a kernel size of 3, followed by layer normalization and a ReLU activation. Residual connections are used throughout the TCN stack, following the design principles of ResNet [16]. In the first TCN block, a pointwise (kernel size = 1) causal temporal convolution is applied to the Prefilter output to match its channel dimension to that of the block output before the residual addition. Operationally, the encoder treats each electrode’s time series as a separate stream by reshaping (𝐵, 𝐸, 𝑇, 1) to (𝐵 × 𝐸, 𝑇, 1) for convolution and restoring the original axes afterward, producing an encoder output 𝑧 ∈ ℝ^𝐵×𝐸×𝑇×𝐹^. Here, 𝐵 denotes the batch size, 𝐸the number of electrodes, 𝑇 the number of temporal samples within each input window, and 𝐹 the number of feature channels produced by the convolutional layers.

**Figure 2.**
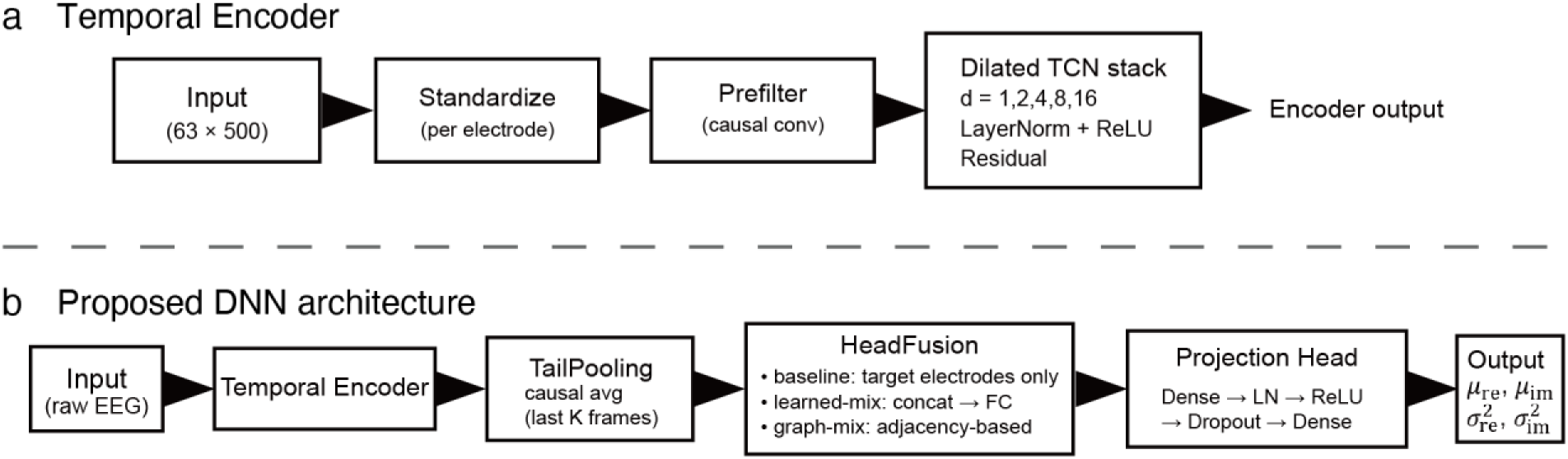
Architecture of the proposed deep neural network. (a) Temporal encoder consisting of per-electrode standardization, a causal prefilter, and a dilated TCN stack with exponentially increasing dilation factors. The encoder produces temporal feature representations for each electrode. (b) Full DNN architecture. Raw multichannel EEG is processed by the temporal encoder, followed by TailPooling, which performs causal averaging over the last 𝐾time frames. HeadFusion integrates per-electrode features using one of three configurations: (i) baseline (target electrodes only), (ii) learned mixing (concatenation followed by a fully connected layer), or (iii) graph-mixing (adjacency-based aggregation). The Projection Head (MLP) outputs the parameters of the predicted analytic signal—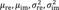 —from which instantaneous phase and amplitude are derived. All models share the same temporal encoder, TailPooling, and Projection Head; only the HeadFusion configuration differs across model variants.

#### 2.2.3 Output Head architecture

The overall model architecture is shown in Figure 2b. Downstream of the temporal encoder, the output head is divided into three consecutive stages—TailPooling, HeadFusion, and Projection.

To summarize recent temporal dynamics, we apply causal averaging over the last K time points (TailPooling). This design is consistent with temporal convolutional models, where predictions primarily rely on the most recent receptive field in causal architectures [11]. Specifically, for the encoder output 𝑧 ∈ ℝ^𝐵×𝐸×𝑇×𝐹^, we compute

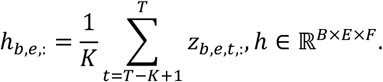

TailPooling thus performs causal averaging along the temporal axis, summarizing short-term temporal dynamics while preserving the feature dimension 𝐹 for each electrode.

The second stage, HeadFusion, aggregates per-electrode features to form a representation suitable for prediction. In the baseline configuration without cross-electrode mixing, only the target electrodes specified by a predefined index set are retained and passed to the next stage. The corresponding per-electrode features are normalized, regularized by dropout, and then flattened before being passed to the following stage. This baseline therefore performs no cross-electrode mixing and relies solely on the target electrodes for prediction. In an alternative learned-mixing configuration, the features from all 63 electrodes are concatenated, and a fully connected layer learns cross-electrode mixing weights directly from the data. Variants that employ missing-signal simulation or graph-based mixing are introduced in Section 2.3.

Finally, the Projection stage maps the fused representation to the analytic-signal parameters through a small multilayer perceptron consisting of a per-electrode dense layer, layer normalization, a ReLU activation, dropout, and a final dense layer. This projection produces the mean values (𝜇_re_, 𝜇_im_) and the corresponding variances 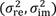 for the real and imaginary components of the analytic signal, which are subsequently used to derive phase and amplitude.

#### 2.2.4 Probabilistic Modeling of Analytic Signals

Exact prediction of EEG signals is often challenging and sometimes impossible because EEG measures brain activity, which is affected by many different factors. Consequently, the training data may include instances where predicting the future from past EEG signals is impossible. Therefore, we modeled the real and imaginary components of the predicted analytic signal as probabilistic outputs and trained the model by maximizing the likelihood of the ground-truth analytic signal. Previous studies have shown that representing prediction uncertainty through probability distributions is effective for DNNs, particularly when the training data contain unpredictable or noisy information [12, 13].

Let

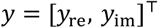

denote the ground-truth analytic signal at the target time point, where 𝑦_re_ and 𝑦_im_ are the real and imaginary components obtained from the Hilbert-transform-based analytic signal, respectively. The model predicts their means (𝜇_re_(𝑥) , 𝜇_im_(𝑥)) and standard deviations (𝜎_re_, 𝜎_im_). The real and imaginary parts of the analytic signal at a specific time point are modeled as a two-dimensional normal distribution:

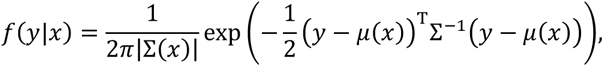

where

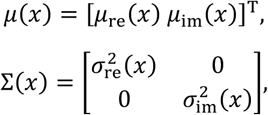

and the diagonal covariance reflects the assumption that the real and imaginary components of the analytic signal are conditionally independent given 𝑥. Here, 𝑥 ∈ ℝ^𝐸×𝑇^ denotes an input sample, and the mean 𝜇(𝑥) and covariance Σ(𝑥) are expressed as functions of 𝑥. For the 𝑛-th sample, we write 𝑥 = 𝑥_𝑛_ and 𝑦 = 𝑦_𝑛_ = [ 𝑦_𝑛,re_, 𝑦_𝑛,im_ ]^⊤^.

The model thus predicts the mean values 𝜇_re_(𝑥_𝑛_) and 𝜇_im_(𝑥_𝑛_) together with the standard deviation values 𝜎_re_(𝑥_𝑛_) and 𝜎_im_(𝑥_𝑛_) for the real and imaginary parts of the analytic signal at a specific time. The estimated values for the real and imaginary parts can then be used to calculate the phase and amplitude. In this study, we specifically focused on predicting the analytic signals at the following time points: −300, −250, −200, −150, −100, −50, 0, 50, 100, 150, 200, 250, and 300 ms. The negative values represent the past, 0 ms represents the present, and the positive values represent the future. We employed the negative log-likelihood (NLL) function of the above normal distribution as the loss function: 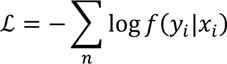

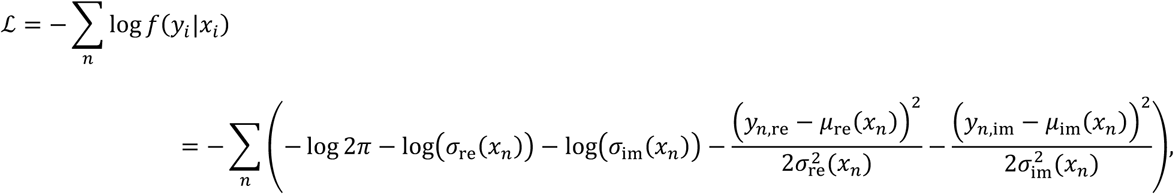

where 𝑦_𝑛,re_, 𝑦_𝑛,im_ is the ground-truth real and imaginary components of 𝑦_𝑛_ corresponding to 𝑥_𝑛_. We trained the model using Adam to minimize the above NLL. For comparison, we also trained a separate point-estimate model using the mean squared error (MSE) loss. This formulation is equivalent to maximizing a Gaussian likelihood with fixed unit variance (𝜎_𝑟𝑒_(𝑥_𝑛_) = 𝜎_𝑖𝑚_(𝑥_𝑛_) = 1), and therefore optimizes only the predicted mean values.

### 2.3 Robustness and Cross-Electrode Integration

This subsection introduces two architectural extensions designed to enhance robustness to missing electrodes and to enable effective cross-electrode information integration: (i) a missing signal imitation (MSI) layer that is active only during training, and (ii) electrode-mixing mechanisms incorporated in the HeadFusion stage of the output head.

#### 2.3.1 Missing Signal Imitation (MSI)

In practical EEG recordings, individual electrodes may occasionally fail and produce implausible signals. To make the model robust to such failures, we introduce a missing-signal imitation (MSI) perturbation that is applied only during training. With missing probability 𝛼, MSI replaces the entire 1-s trace of electrode 𝑖 in sample 𝑛 with an artificially corrupted signal:

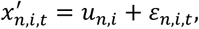

where 𝑢_𝑛,𝑖_ ∼ 𝒰(−500,500) is an electrode-wise baseline value and 𝜀_𝑛,𝑖,𝑡_ ∼ 𝒩(0,1) is small Gaussian noise. Thus, masked electrodes exhibit approximately constant but clearly non-physiological activity over time, mimicking broken or saturated channels that the network should learn to ignore. This stochastic masking discourages the model from over-relying on any single electrode and increases robustness to missing channels. The mechanism is conceptually similar to Dropout, which randomly suppresses units during training to improve generalization [17]. It also empirically improves the calibration of predictive uncertainty for models trained under the NLL objective, consistent with prior work showing that probabilistic deep networks can benefit from variance-aware learning [13].

#### 2.3.2 Electrode Mixing in the Head

Two alternative approaches are used to integrate information across electrodes in the HeadFusion stage. These approaches differ in whether the model learns electrode relationships directly from data or incorporates an explicit adjacency structure derived from the electrode layout.

##### Learned Electrode Mixing (All-Flatten → Dense)

The per-electrode features ℎ ∈ ℝ^𝐵×𝐸×𝐹^ are concatenated and flattened into a vector of size 𝐸 × 𝐹. A fully connected layer then learns cross-electrode mixing weights directly from the data. This formulation makes no explicit assumption about spatial adjacency, allowing the network to infer useful electrode combinations, similar in spirit to data-driven spatial integration used in compact EEG architectures such as EEGNet [14]. In the ablation studies, this configuration is evaluated both with and without MSI applied during training.

##### Graph α-Mix (Adjacency-Guided Aggregation)

When a binary adjacency matrix 𝐴 ∈ {0,1}^𝐸×𝐸^ is available (for example, from the physical electrode layout), we compute a graph-mixed representation by multiplying 𝐴 with the per-electrode features ℎ, i.e., 𝐴ℎ. This design introduces a simple spatial inductive bias inspired by classical EEG spatial filtering approaches, including current source density (CSD) and related Laplacian-based methods, which emphasize local neighborhood structure and are commonly used to reduce the impact of volume conduction effects [18]. The matrix 𝐴 is row-normalized and can be augmented with a self-loop on the diagonal to retain self-information. The mixed representation is then blended with the original ℎ through a learnable scalar gate 𝛼 ∈ [0,1]:

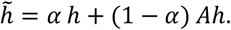

Here 𝐴 is fixed and not optimized, whereas 𝛼 is trained jointly with the rest of the network to determine the contribution of adjacency-based aggregation. In the present study, adjacency-guided mixing is applied uniformly to all electr

#### 2.3.3 Training and Model Variants

Unless otherwise noted, probabilistic modeling and optimization follow Section 2.2, using the negative log-likelihood (NLL) as the primary objective and the mean squared error (MSE) as a point-estimate baseline. The MSI perturbation is applied only during training. Electrode-mixing mechanisms modify only the HeadFusion stage; the temporal encoder and loss functions remain unchanged.

In total, four model configurations are examined (table 1):

1. DNN-NLL: HeadFusion uses target-only selection and flattening (no cross-electrode mixing).
2. DNN-MSE: same as the baseline configuration above but trained with the MSE objective.
3. DNN-NLL + MSI: includes the MSI layer and uses learned cross-electrode mixing (All-Flatten → Dense).
4. DNN-NLL + MSI + Graph: combines MSI with the Graph 𝛼-Mix, in which the adjacency matrix 𝐴 is fixed and only the scalar gate 𝛼 is learned.

**Table 1:** Summary of the model configurations evaluated. All models share the same temporal encoder and TailPooling → HeadFusion → Projection pipeline. The models differ only in their loss functions (NLL or MSE) and the presence of MSI or adjacency-guided fusion.

| Model name | Loss function | MSI (training-time masking) | Electrode mixing (HeadFusion) | Graph-based mixing | Notes |
| --- | --- | --- | --- | --- | --- |
| DNN-NLL | NLL | – | Gather→Flatten (target electrodes only) | – | Baseline model |
| DNN-MSE | MSE | – | Gather → Flatten | – | Point-estimate baseline |
| DNN-NLL+ MSI | NLL | ✓ | All-Flatten→Dense (learned mixing) | – | Tests effect of MSI and learned cross-electrode mixing |
| DNN-NLL+ MSI + Graph | NLL | ✓ | α-controlled fusion | ✓<br>(fixed adjacency A) | Tests adjacency-guided mixing + MSI |

These four models share the same temporal encoder and TailPooling → HeadFusion → Projection pipeline, differing only in the loss function (NLL or MSE) and in whether MSI and graph-based electrode mixing are employed.

### 2.4 EEG data and Validation methods

We validated the method through offline analyses using open EEG data collected from 25 participants (s001–s025) at rest with their eyes closed [19]. The recordings were obtained from 63 EEG electrodes placed according to the international 10/10 system, with AFz as the ground electrode and the left earlobe serving as the reference. EEG signals were sampled at 1000 Hz and band-limited online using low- and high-cutoff frequencies of 0.016 and 250 Hz, respectively. After recording, the reference was rereferenced to the average of the right and left earlobes, and the sampling rate was down-sampled to 500 Hz.

In this study, we adopted two evaluation protocols using this dataset: a subject-wise evaluation and a cross-subject evaluation. In the subject-wise protocol, each participant’s EEG data were independently split into training, validation, and test sets, and the model was trained and evaluated separately for each participant. The EEG time series were divided into 14 consecutive segments; segments 1–10 were used for training, segments 11–12 for validation, and segments 13–14 for testing. Notably, the validation and test segments always contained data occurring after the training segments to avoid temporal leakage. Using this protocol, we trained and compared the performance of the DNN-NLL and DNN-MSI configurations for participants 1–10. We also investigated the effect of model depth by increasing the number of layers in the Dilated TCN stack from 3 to 5. Evaluation metrics included the mean squared error (MSE) for continuous prediction of EEG amplitude and phase, and classification accuracy for discrete phase-prediction tasks. Phase classification was performed by assigning predicted phases to the closest 45° bin and computing accuracy. MSE quantifies the magnitude of prediction errors, whereas accuracy measures the proportion of correctly predicted phase labels within a defined tolerance. These metrics were computed for both network depths and across all ten participants to provide a comprehensive comparison of architectural and training-objective choices.

In addition to the subject-wise evaluation, we performed a cross-subject validation to assess generalization to previously unseen individuals—a requirement for practical real-time EEG phase prediction. For this analysis, the dataset of 25 participants was partitioned into 10 training subjects and 15 test subjects, each further subdivided into training, validation, and test segments using the same temporal split described above. Model performance was then assessed on the test segments of subjects entirely unseen during training. This cross-subject evaluation demonstrates the model’s ability to generalize to new individuals and highlights its suitability for real-world deployment.

For the ground-truth analytic signal, we selected five posterior electrodes (PO7, PO3, POz, PO4, and PO8) within the 8–13 Hz frequency band. The analytic signal for each participant was computed using a band-pass filter followed by the Hilbert transform. Both past and future EEG samples were used to compute the analytic signal, which served as the ground-truth target during model training. Each dataset therefore comprised: (i) 1 s of raw EEG signals from all 63 electrodes as model input, (ii) an adjacency matrix representing electrode connectivity, and (iii) the analytic-signal ground-truth data described above.

### 2.5 Missing-channel evaluation

In addition to the above analyses, we conducted a missing-channel evaluation to assess model robustness when critical electrodes fail. In this experiment, we focused on the five posterior electrodes used as prediction targets (PO7, PO3, POz, PO4, and PO8). For each test sample and for each of these electrodes in turn, the corresponding input channel was artificially set to a missing state, and the model was required to predict the analytic signal of that very electrode. This setting reflects practical scenarios in which a sensor becomes detached during real-time EEG acquisition. The evaluation was performed under the cross-subject protocol, ensuring that missing-channel robustness was assessed on subjects unseen during training. We compared three configurations: DNN-NLL, DNN-NLL+MSI, and DNN-NLL+MSI+Graph α, thereby evaluating the contribution of MSI-based perturbation and adjacency-guided mixing to the recovery of missing channels.

## 3. Results

### 3.1 Prediction Performance with subject-wise evaluation

The AR baseline used here followed the settings described in Section 2.1. Figure 3 summarizes the prediction performance under the subject-wise evaluation. Both DNN models outperformed the AR baseline across all metrics around the current time point. In terms of phase accuracy, the AR baseline showed a rapid decline near 0 ms, whereas the DNN-MSE and DNN-NLL models maintained substantially higher accuracy. The DNN-NLL model achieved the best performance overall, exhibiting the highest accuracy and the lowest phase and amplitude MSEs at 0 ms. Similar trends were observed in the MSE plots, where the AR baseline showed large errors, particularly for future prediction windows, while the DNN models demonstrated gradual and smaller increases in error.

**Figure 3:**
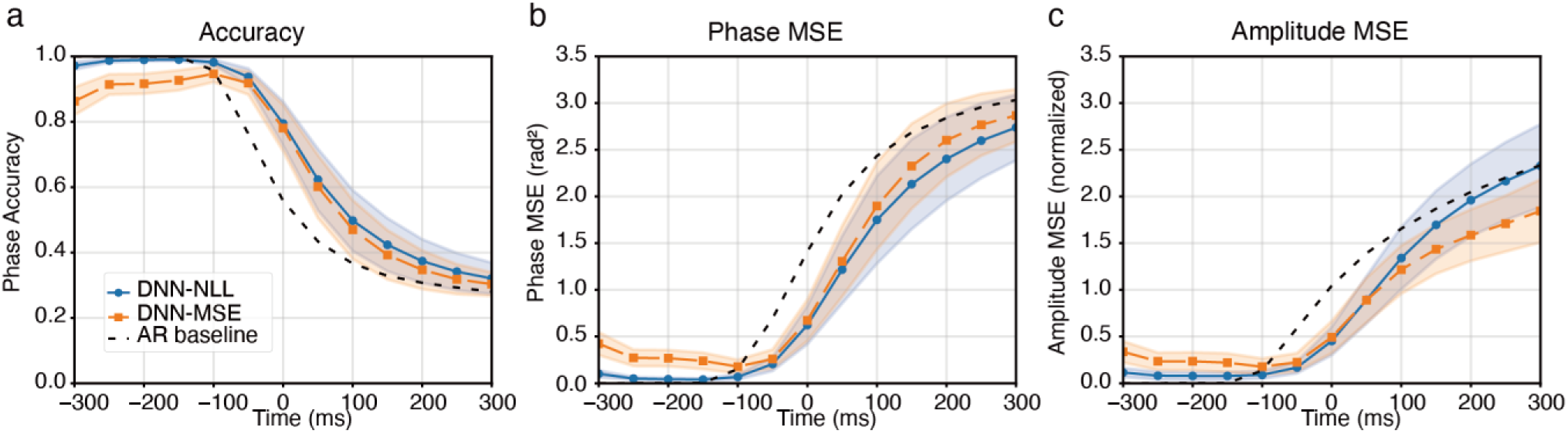
Prediction performance under the subject-wise evaluation. (a) Phase accuracy, (b) phase MSE, and (c) amplitude MSE for the AR baseline (black dashed), DNN-NLL (blue), and DNN-MSE (orange), all configured with a stack size of three. Shaded regions denote the standard deviation across subjects. The DNN-NLL model shows the best overall performance, particularly around 0 ms and at future time points, while the AR baseline performs relatively well only for past time points. These results highlight the advantage of probabilistic training for stable phase and amplitude prediction across subjects.

Table 2 provides a quantitative comparison of the prediction performance at 0 ms for different stack sizes. Among the tested configurations, the three-stack DNN-NLL model consistently achieved strong performance across evaluation metrics, without clear gains from additional depth. Increasing the stack size beyond three resulted in only marginal improvements or slight degradation, indicating diminishing returns in model depth. These results demonstrate that the proposed DNN-NLL architecture can robustly estimate analytic signals from raw EEG data and that a three-stack configuration provides an optimal balance between model complexity and predictive performance.

**Table 2:** Prediction performance at 0 ms under the subject-wise evaluation for the AR baseline, DNN-NLL, and DNN-MSE models with different stack sizes. Values represent the mean ± standard deviation across subjects. The DNN-NLL model with three stacks achieved the best overall performance.

| Method | Stack($n_{blocks}$) | accuracy | MSE(Phase) | MSE(Amplitude) | N |
| --- | --- | --- | --- | --- | --- |
| AR prediction | - | $0.5581 \pm 0.0382$ | $1.4110 \pm 0.1684$ | $1.0453 \pm 0.0863$ | 10 |
| DNN-NLL | 3 | $0.7946 \pm 0.0299$ | $0.6239 \pm 0.0910$ | $0.4528 \pm 0.0672$ | 10 |
| | 4 | $0.7926 \pm 0.0303$ | $0.6369 \pm 0.0939$ | $0.4666 \pm 0.0769$ | 10 |
| | 5 | $0.7920 \pm 0.0304$ | $0.6314 \pm 0.0930$ | $0.4616 \pm 0.0719$ | 10 |
| DNN-MSE | 3 | $0.7809 \pm 0.0314$ | $0.6742 \pm 0.0980$ | $0.4944 \pm 0.0758$ | 10 |
| | 4 | $0.7676 \pm 0.0327$ | $0.7168 \pm 0.1052$ | $0.5106 \pm 0.0732$ | 10 |
| | 5 | $0.7601 \pm 0.0321$ | $0.7412 \pm 0.1037$ | $0.5540 \pm 0.0872$ | 10 |

### 3.2 Prediction Performance with cross-subject evaluation

Figure 4 shows the prediction performance in the cross-subject evaluation, where models were trained on data from multiple subjects and tested on an unseen subject. Compared with the AR baseline, all TCN-based models demonstrated substantially higher phase accuracy and lower MSE across all time points. Around 0 ms, the DNN-NLL model achieved the best performance, followed closely by the DNN-NLL+MSI and DNN-NLL+MSI+Graph models.

**Figure 4:**
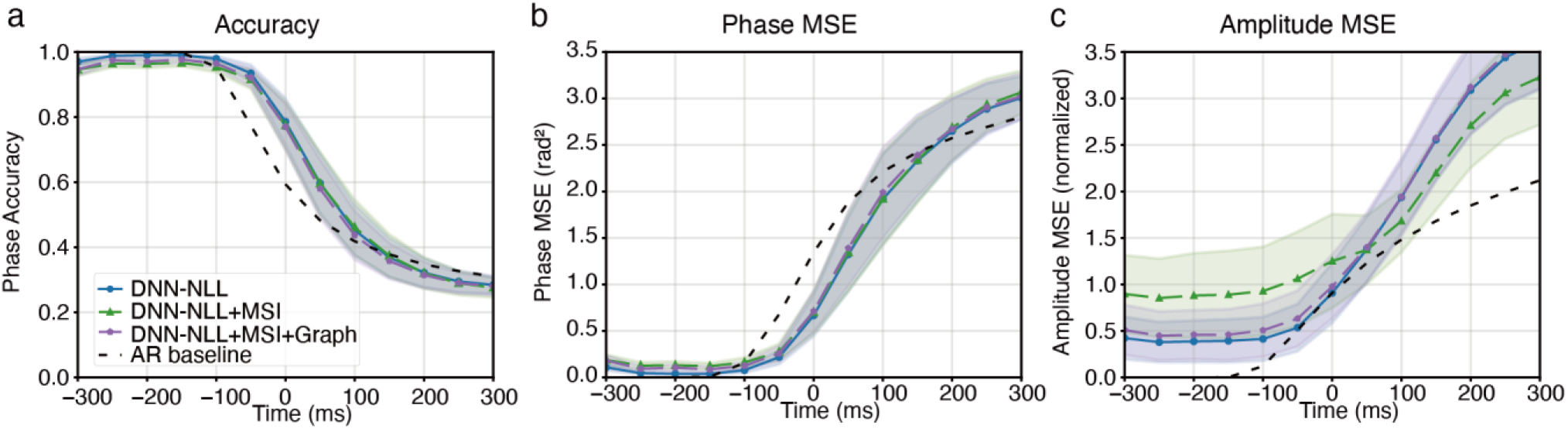
Prediction performance under the cross-subject evaluation. (a) Phase accuracy, (b) phase MSE, and (c) amplitude MSE for DNN-NLL (blue), DNN-NLL+MSI (green, 𝑝 = 0.2), DNN-NLL+MSI+Graph (purple, 𝑝 = 0.2), and the AR baseline (black dotted). Shaded regions indicate the standard deviation across subjects. All DNN models outperformed the AR baseline across most time points, particularly around 0 ms. The probabilistic models (DNN-NLL variants) maintained stable performance in both past and future intervals, with MSI and GraphMix providing additional robustness.

Although the introduction of MSI slightly increased prediction errors in some cases, both MSI variants maintained performance levels comparable to DNN-NLL and clearly outperformed the AR-based method. These results indicate that the proposed TCN architecture generalizes well across subjects and preserves its predictive ability even when trained on heterogeneous EEG data.

### 3.3 Prediction Performance with Missing-channel Evaluation

Figure 5 presents the prediction performance when a subset of electrodes was replaced with missing values. Under this challenging setting, the baseline DNN-NLL model exhibited a noticeable drop in phase accuracy, particularly around 0 ms and in future time points. In contrast, both MSI-based models substantially mitigated the performance degradation caused by missing channels. Among them, the DNN-NLL+MSI+Graph model demonstrated the highest robustness, maintaining accuracy levels close to those observed under complete-channel conditions.

**Figure 5:**
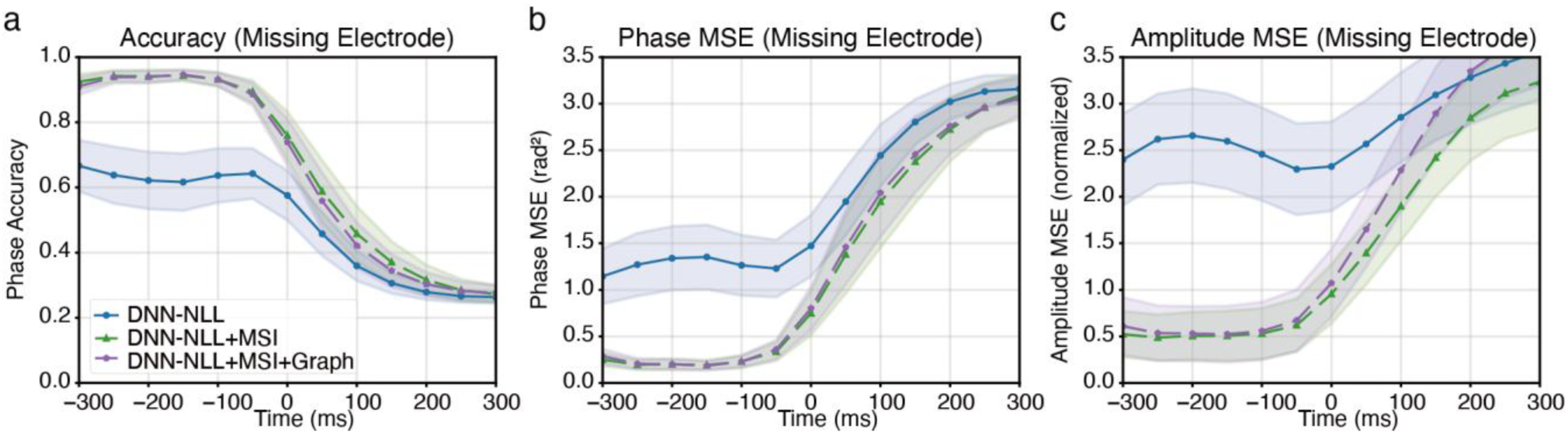
Prediction performance under missing-channel conditions in the cross-subject evaluation. (a) Phase accuracy, (b) phase MSE, and (c) amplitude MSE for DNN-NLL (blue), DNN-NLL+MSI (green, 𝑝 = 0.2), and DNN-NLL+MSI+Graph (purple, 𝑝 = 0.2). Shaded areas represent the standard deviation across subjects. Missing channels notably degraded the performance of the baseline DNN-NLL model, whereas MSI-based variants preserved high accuracy and low error across time. Among them, DNN-NLL+MSI+Graph exhibited the greatest robustness, showing minimal sensitivity to missing electrodes.

These results indicate that MSI-based training provides substantial benefits in scenarios where electrode failures or noise contamination occur during EEG acquisition by reducing over-reliance on individual channels. In all subsequent experiments, we set the missing-signal probability to 𝑝 = 0.2, which was selected based on preliminary analyses as a reasonable compromise that introduces sufficient channel perturbation during training without noticeably degrading performance under normal conditions.

### 3.4 Normalized Residual Distributions of the Predicted Signals

To evaluate the calibration of the predicted variance, we analyzed normalized residuals computed on the test data in the cross-subject setting. This evaluation focuses on generalization to unseen subjects rather than performance on the training data. The normalized residuals were defined as

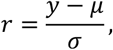

where 𝑦 denotes the ground-truth analytic signal and 𝜇 and 𝜎 are the predicted mean and standard deviation. The normalized residuals were computed separately for the real and imaginary components and then pooled for analysis. For a well-calibrated model, the distribution of 𝑟 should follow a standard normal distribution.

Figure 6 shows the normalized residual distributions for the NLL, MSI (𝑝 = 0.2), and MSI+Graph (𝑝 = 0.2) models under both normal and missing-electrode conditions. The NLL model exhibits substantial overdispersion, especially when an electrode is missing, indicating that its predicted variance severely underestimates the true prediction error. In contrast, both MSI and MSI+Graph models produce residual distributions that more closely match 𝒩(0,1), demonstrating improved calibration of the predicted variance. The effect is particularly pronounced under missing-electrode conditions, where MSI-based models greatly reduce overdispersion. These results confirm that the MSI mechanism not only enhances robustness to missing channels but also leads to more reliable predictive uncertainty.

**Figure 6:**
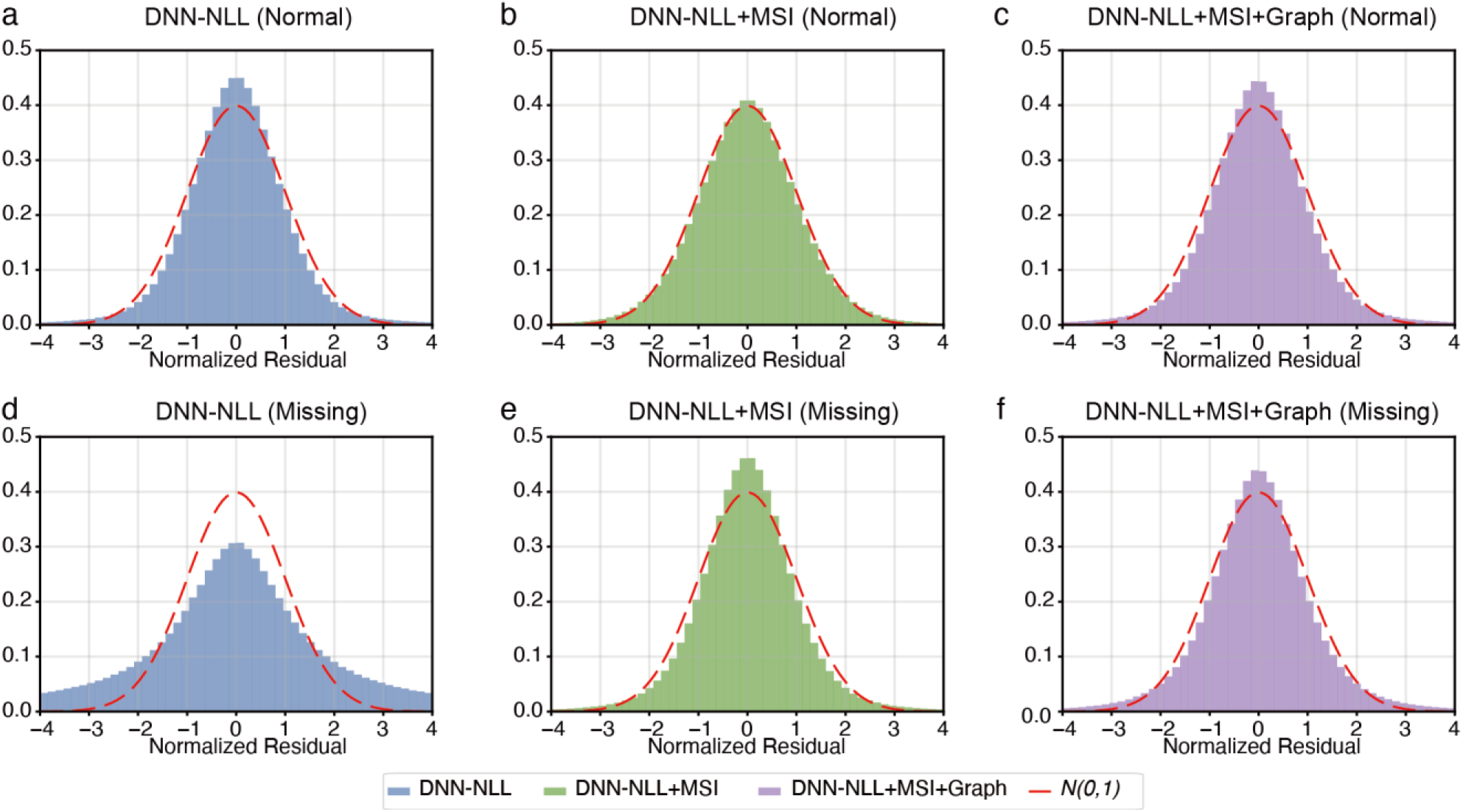
Normalized residual histograms for the NLL, MSI (𝑝 = 0.2), and MSI+Graph (𝑝 = 0.2) models under normal and missing-electrode conditions. (a–c) Histograms of normalized residuals under the normal condition for (a) DNN-NLL, (b) DNN-NLL+MSI, and (c) DNN-NLL+MSI+Graph. (d–f) The corresponding histograms under the missing-electrode condition for DNN-NLL (d), DNN-NLL+MSI (e), and DNN-NLL+MSI+Graph (f). Each histogram is compared with the standard normal distribution *𝑁*(0,1) (red dashed line). The NLL model exhibits clear overdispersion—especially under missing-electrode conditions—indicating underestimation of predictive variance. In contrast, both MSI-based models yield residual distributions that more closely match *𝑁*(0,1), demonstrating improved variance calibration and greater robustness to sensor dropout.

## 4. Discussion

Deep learning has been widely applied to multichannel EEG analysis [7,8]. Although many studies explicitly aim at cross-subject EEG modeling, performance tends to degrade when models are applied to unseen subjects, reflecting substantial inter-subject variability. In this study, we investigate cross-subject generalization in the context of direct analytic signal prediction from raw multichannel EEG. The principal contribution of this work is the demonstration that a single model trained on data from multiple participants can achieve robust cross-subject prediction and consistently outperform a conventional AR-based method across evaluation metrics. These results suggest that the proposed temporal convolutional architecture can reliably approximate band-limited analytic signal representations across subjects, despite variability in signal amplitude, electrode impedance, and spatial distribution of neural generators.

Results from both subject-wise and cross-subject evaluations highlight the versatility of the proposed architecture (Figures 3 and 4). In the subject-wise setting, the DNN outperformed the AR baseline around the current time point and in future prediction windows for both phase and amplitude (Figure 3). In the cross-subject setting, robust phase prediction performance was preserved without subject-specific retraining, whereas amplitude prediction exhibited more limited generalization across subjects (Figure 4). These results indicate that the proposed framework supports stable cross-subject phase estimation, while amplitude prediction remains more sensitive to inter-individual differences.

The differing cross-subject behavior between phase and amplitude predictions can be partly explained by their distinct signal characteristics. Instantaneous phase is a circular and scale-invariant variable, making it relatively robust to inter-individual differences such as electrode impedance, head geometry, and overall signal scaling. In contrast, amplitude is a non-circular and absolute quantity that directly reflects signal magnitude, and is therefore more sensitive to subject-specific factors and recording conditions. As a result, amplitude prediction poses a greater challenge in cross-subject settings, even when the underlying oscillatory structure is shared across individuals.

A central finding of this work concerns the role of the missing-signal imitation (MSI) mechanism in improving robustness under realistic recording conditions. MSI was originally introduced to address electrode dropout, a common occurrence in practical EEG acquisition. The results demonstrate that MSI-based models preserve prediction performance even when critical electrodes are removed during testing, whereas models trained without MSI exhibit substantial degradation. This robustness is particularly important for long-duration or mobile EEG recordings, where sensor quality and channel availability may fluctuate over time.

Beyond robustness to missing electrodes, MSI also acts as an effective regularizer during training, mitigating overfitting and stabilizing model behavior across subjects, similar to Dropout [17]. Models trained with a moderate missing probability (𝑝 = 0.2) achieved a favorable balance between performance under normal conditions and resilience to channel loss. Importantly, MSI further contributed to improved calibration of predictive uncertainty. Residual analyses (Figure 6) revealed that models trained without MSI systematically underestimated prediction variance, resulting in over-dispersed standardized residuals, whereas MSI-based training produced residual distributions more closely aligned with the standard normal distribution [12,13]. These findings indicate that MSI enhances not only robustness to incomplete observations but also the reliability of probabilistic predictions, which is essential for practical deployment of EEG-based systems.

The electrode-mixing strategies further contributed to model robustness. Learned mixing enabled flexible integration of spatial information across electrodes, whereas the graph-based approach leveraged adjacency relationships to guide feature propagation. The graph-based mixing did not consistently outperform learned mixing, but showed comparable robustness under missing-electrode conditions (Figure 5). More broadly, classical EEG spatial filtering techniques—including current source density, surface Laplacian, and related local derivations—share the common objective of reducing volume conduction by exploiting electrode geometry and spatial locality [20]. These results indicate that while adjacency-based mixing can be applied without severe degradation, its benefits over fully learned mixing remain limited in the present setting. In the present framework, electrode mixing plays two complementary roles: enhancing robustness to missing electrodes and introducing a spatial inductive bias that promotes local integration across neighboring sensors. The former is particularly important under realistic recording conditions, while the latter is conceptually related to classical EEG spatial filtering approaches that aim to mitigate volume conduction effects [20].

Previous studies, including our Kalman-filter-based real-time phase estimation framework [21], demonstrated that state-space modeling can stabilize instantaneous phase estimation without requiring data-driven learning. Notably, the EEG recordings used in that study are included within the dataset analyzed in the present work. However, Kalman filtering differs fundamentally from the proposed DNN approach in both its underlying model assumptions and its definition of instantaneous phase. Kalman- and AR-based methods rely on linear state-space formulations and predefined bandpass filters, and do not exploit learned predictive structure shared across subjects. Instead, their parameters are estimated online in real time for each recording, without requiring prior training on data from multiple participants. Moreover, the phase extracted in [21] is not directly comparable to the analytic-signal-based phase used in the present study, making quantitative comparison inappropriate despite the overlap in data. For these reasons, we report comparisons only with the AR baseline, which shares the same analytic-signal formulation as the proposed method and provides a methodologically consistent reference point.

Although the present study focuses on offline analyses, the architecture offers promising foundations for future real-time systems. In particular, the demonstrated cross-subject generalization reduces the need for per-user calibration, and the robustness to missing channels supports stable operation under realistic recording conditions. Future work will explore the computational and architectural modifications required for low-latency deployment, as well as extensions to task-related or stimulation-locked EEG paradigms.

### Limitation

This work has several limitations. First, the proposed model requires offline training on a substantial dataset and was not evaluated under strict real-time constraints; additional engineering will be necessary to guarantee low-latency deployment on specific hardware. Second, we considered a single EEG recording setup and frequency band, and it remains to be tested how well the model transfers to different montages, frequency ranges, and noise conditions. Third, the graph structure used for adjacency-based mixing was derived from the physical electrode layout and may not fully capture functional connectivity. Future work could explore data-driven or individualized graph structures and investigate extensions to task-related and stimulation-locked EEG paradigms.

In conclusion, this study demonstrates that deep learning enables accurate and generalizable prediction of instantaneous EEG state across individuals, surpassing conventional methods in accuracy, robustness, and uncertainty estimation. By reducing dependence on manual parameter selection and enhancing tolerance to sensor variability, the proposed framework provides a scalable foundation for next-generation EEG applications, including automated experimental paradigms, large-cohort neural monitoring, and future closed-loop systems.

## 5. Conclusion

This study presented a deep learning framework for direct prediction of analytic EEG signals from raw multichannel input, emphasizing its ability to generalize across individuals. The proposed model outperformed conventional AR-based method in both subject-wise and cross-subject evaluations and demonstrated robustness to missing channels through the incorporation of missing-signal imitation and electrode-mixing strategies. Furthermore, the probabilistic formulation and MSI-driven calibration improvements enabled more reliable estimation of prediction uncertainty.

By reducing dependence on manual parameter selection and achieving stable performance across heterogeneous participants, the proposed approach provides a scalable foundation for future EEG applications. Although the present study focused on offline analyses, the findings support the potential for extending this framework toward real-time implementations and closed-loop experimental paradigms.

## Declaration of generative AI and AI-assisted technologies in the manuscript preparation process

During the preparation of this work, the authors used ChatGPT (OpenAI, GPT-5.1) to improve the clarity and grammar of English text. After using this tool, the authors reviewed and edited the content as needed and take full responsibility for the content of this publication.

## Acknowledgements

We gratefully acknowledge the use of publicly available open EEG datasets, which were instrumental in the development and validation of this research. We extend our appreciation to the researchers and institutions who made these datasets accessible to the scientific community. This work was supported by JSPS KAKENHI Grant Numbers 20K19887 and 24K18601. T.O. holds a visiting appointment at the Institute of Statistical Mathematics.

